# pQEB1: a hospital outbreak plasmid lineage carrying *bla*_KPC-2_

**DOI:** 10.1101/2024.06.07.597914

**Authors:** Robert A. Moran, Mahboobeh Behruznia, Elisabeth Holden, Mark I. Garvey, Alan McNally

**Author notes:** Correspondence: Alan McNally.

## Abstract

While conducting genomic surveillance of carbapenemase-producing Enterobacteriaceae (CPEs) from patient colonisation and clinical infections at Birmingham’s Queen Elizabeth Hospital (QE), we identified an N-type plasmid lineage, pQEB1, carrying several antibiotic resistance genes including the carbapenemase gene *bla*_KPC-2_. The pQEB1 lineage is concerning due to its conferral of multi-drug resistance, its host range and apparent transmissibility, and its potential for acquiring further resistance genes. Representatives of pQEB1 were found in three sequence types (STs) of *Citrobacter freundii*, two STs of *Enterobacter cloacae*, and three species of *Klebsiella*. Hosts of pQEB1 were isolated from 11 different patients who stayed in various wards throughout the hospital complex over a 13-month period from January 2023 to February 2024. At present, the only representatives of the pQEB1 lineage in GenBank were carried by an *Enterobacter hormaechei* isolated from a blood sample at the QE in 2016 and a *Klebsiella pneumoniae* isolated from a urine sample at University Hospitals Coventry and Warwickshire (UHCW) in May 2023. The UHCW patient had been treated at the QE.

Long-read whole-genome sequencing was performed on Oxford Nanopore R10.4.1 flow cells, facilitating comparison of complete plasmid sequences. We identified structural variants of pQEB1 and defined the molecular events responsible for them. These have included IS*26*-mediated inversions and acquisitions of multiple insertion sequences and transposons, including carriers of mercury or arsenic resistance genes. We found that a particular inversion variant of pQEB1 was strongly associated with the QE Liver speciality after appearing in November 2023, but was found in different specialities and wards in January/February 2024. That variant has so far been seen in five different bacterial hosts from six patients, consistent with recent and ongoing inter-host and inter-patient transmission of pQEB1 in this hospital setting.

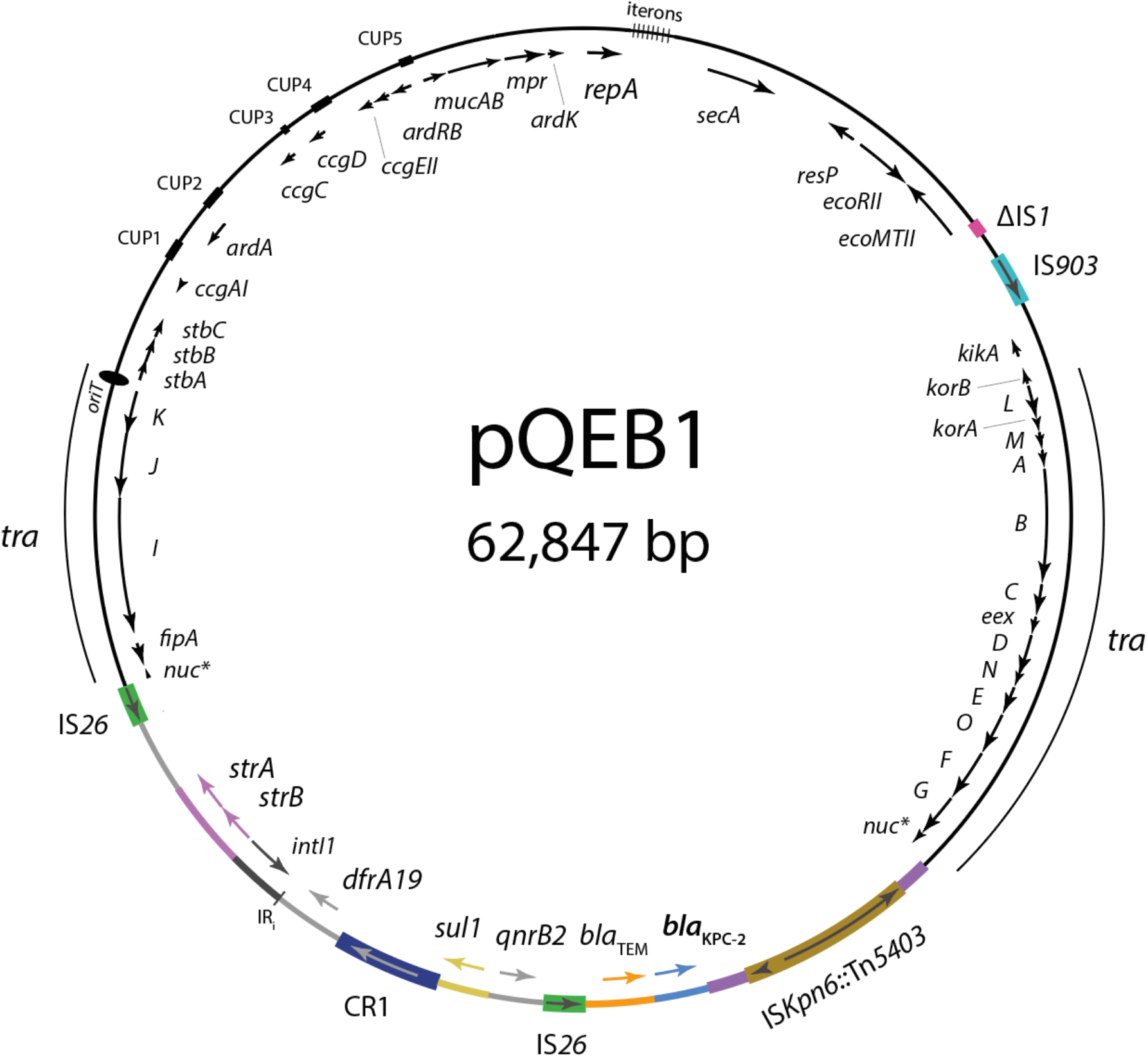

## Introduction

University Hospitals Birmingham NHS Foundation Trust (UHB) is one of the largest UK hospital trusts, spanning four sites that cover the majority of the Birmingham population. The Queen Elizabeth (QE) site is a 1200-bed tertiary referral centre that includes a 100-bed intensive care unit (ICU), which is the largest co-located ICU globally. The QE is a level 1 trauma centre, houses the largest solid organ transplant service in Europe, and sees >400 repatriated patients per year who have received healthcare abroad. As a specialist tertiary centre, the use of carbapenem antibiotics at the QE is one of the highest in England ^1^. To date, QE clinical laboratories have isolated over 450 carbapenemase-producing Enterobacteriaceae (CPE), with approximately 66% of these producing metallo-ß-lactamases KPC or NDM. Around 7% of CPEs at the QE have been isolated from bloodstream infections, and all-cause mortality in this patient group is 70% within a year.

Plasmids are associated with carbapenem resistance in Enterobacteriaceae globally. While plasmids drive the dissemination of carbapenem resistance genes (CRGs) between bacterial hosts, smaller mobile genetic elements (MGEs) such as insertion sequences (ISs) and transposons drive CRG dispersal between plasmids, or between plasmids and host chromosomes ^2^. Plasmid topologies, or structures, are shaped by the actions of these smaller MGEs, which can insert into plasmids, delete or invert parts of them, and combine distinct plasmid backbones with one another in the form of cointegrates ^3–5^. It is crucial to study plasmid structures in order to understand their evolution. Informed comparative analyses can also provide insights into the ongoing epidemiology of plasmids in clinical settings. An understanding of how plasmid structures shift following diverse molecular events permits the integration of non-identical plasmid variants in transmission assessments for evolving lineages.

In order to develop a better understanding of the bacterial strains and plasmids associated with CPE colonisation and infections at the QE, we have been monitoring carbapenem-resistant isolates by performing whole-genome sequencing using Oxford Nanopore R10.4.1 flow cells. With the latest flow cell chemistry, this sequencing method enables cost and time-efficient genomic surveillance of bacterial pathogens and the mobile genetic elements associated with antibiotic resistance ^6^. We report our findings here to illustrate the utility of the approach in this setting, and to highlight the insights our study has so far provided into the biology of a carbapenem resistance plasmid involved in what appears to be an ongoing outbreak.

## Results

The circular plasmid pQEB1 was found in the complete genome of a ST527 *E. cloacae* that was isolated from a patient blood sample at the QE in January 2023. pQEB1 is 62,847 bp and largely comprised of a backbone related to that of the reference IncN plasmid R46 (GenBank accession AY046076). The pQEB1 backbone is interrupted by an IS*903*-like element, a partial copy of IS*1*, and a complex 19,918 bp antibiotic resistance region (Figure 1A). The resistance region interrupts the *nuc* gene and is bounded by IS*Kpn6*::Tn*5403* at one end and IS*26* at the other. It includes complete or partial sequences of at least five further mobile genetic elements: Tn*2*, Tn*4401*, Tn*5393*, CR1 and at least one class 1 integron. The *bla*_KPC-2_ carbapenemase gene is in a fragment of Tn*4401*, and the resistance region also contains *bla*_TEM_, the sulphonamide resistance gene *sul1*, trimethoprim resistance gene *dfrA19*, streptomycin resistance genes *strAB*, and quinolone resistance gene *qnrB2*.

**Figure 1:**
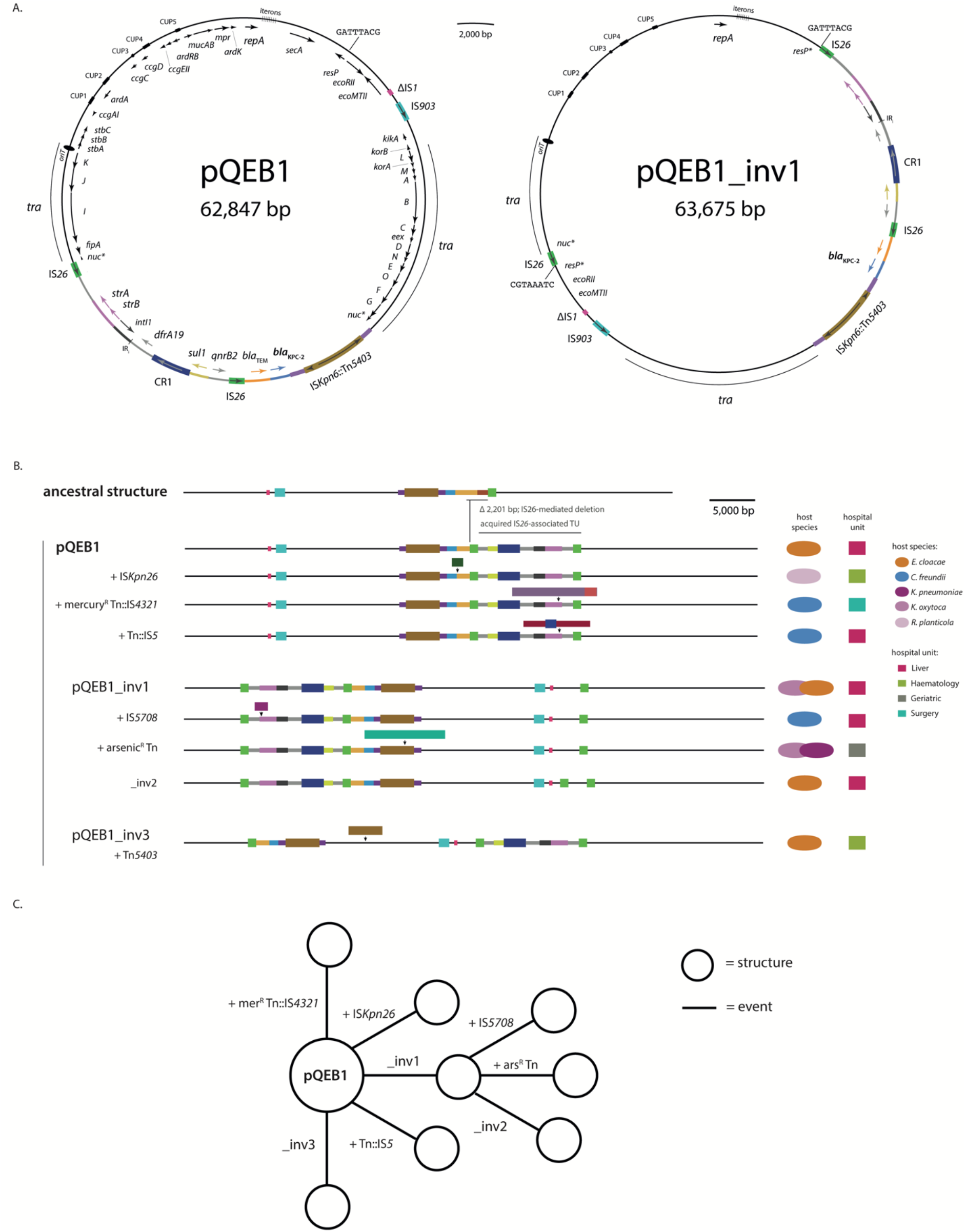
*bla*_KPC-2_-bearing plasmid lineage pQEB1. A) Circular maps of pQEB1 and inversion variant pQEB1_inv1. Gene names are labelled on the inside of circles, and mobile genetic element names on the outside. The extents of transfer (*tra*) regions are marked with labelled arcs and the positions of iteron sequences are indicated. B) Scaled, schematic maps of pQEB1 variants observed here, beneath the putative ancestral structure, “IncN TypeB”. Each structure is labelled with the molecular event that generated it. Host species and locations of isolation are indicated to the right. C) Stepwise evolution of pQEB1 variants from the reference sequence. Structures shown in part B are represented by circles, and the evolutionary events that generated them by labelled lines.

pQEB1 appears to be derived from a plasmid, previously described and named “IncN TypeB” (TypeB), that was carried by a *K. pneumoniae* isolated in the UK in or prior to 2016 ^7^. The resistance regions in pQEB1 and TypeB are inserted at precisely the same backbone position, but in pQEB1 the region has lost 2,201 bp in an IS*26*-mediated deletion event and contains a 10,539 bp IS*26*-flanked segment where TypeB contains a single IS*26* (Figure 1B). The presence of the 10,539 bp segment in pQEB1 can be explained by a targeted conservative IS*26* transposition event, which would be expected to produce a structure like this following integration of a translocatable unit (TU) at the single IS*26* in TypeB ^8^. A plasmid closely related to pQEB1 has recently been described in association with an outbreak of KPC-2 in Germany ^9^. The plasmid circulating in Germany has the same backbone as TypeB and pQEB1, with a resistance region inserted at the same position, including the same ARG-bearing IS*26* TU as pQEB1. However, structural differences within its resistance region clearly distinguish the German lineage (represented by GenBank accession CP104944) from pQEB1. For example, although both feature regions bounded at one end by IS*Kpn6*, in pQEB1 IS*Kpn6* has been interrupted by Tn*5403*, and the German lineage includes an additional 11,821 bp IS*26*-flanked segment. We therefore define the pQEB1 lineage on the basis of the structure of its backbone and resistance region shown in Figure 1A. We used the complete sequence of pQEB1 to query GenBank (last search May 15, 2024), which returned only two representatives of the lineage. These were an ancestral plasmid found in an *E. hormaechei* isolated from a blood sample at the QE in 2016 (CP035387), and a plasmid with identical structure to pQEB1 in a *K. pneumoniae* isolated from a urine sample at UHCW in May 2023 (CP141849) ^10^.

We found representatives of pQEB1 in 11 further CPEs isolated from 10 different QE patients between February 2023 and February 2024 (Table 1). Hosts of pQEB1 included three sequence types (STs) of *Citrobacter freundii*, two of *Enterobacter cloacae*, and three species of *Klebsiella*: *pneumoniae, oxytoca* and *planticola* (Table 1). These were isolated in seven different hospital wards, including in a Geriatric ward that is located in a separate building to the one that hosts the rest of the specialties listed in Table 1. Representatives of pQEB1 ranged from 62,847 bp to 72,485 bp in size and exhibited nine different structures (Figure 1B, Table 1). We identified structural differences relative to our reference sequence, and determined the molecular events that gave rise to each structure found here, revealing that the evolution of pQEB1 has been shaped by the actions of various mobile genetic elements (Table 1). pQEB1 variants have been impacted by three different inversion events mediated by IS*26*, and insertion events involving four different transposons and four different insertion sequences (Table 1). Notable amongst the acquired elements were transposons carrying arsenic or mercury resistance genes. In four of six cases, copies of the newly-acquired elements in pQEB1 variants were found in either the chromosome or co-resident plasmids of their current hosts (Table 1).

**Table 1:**
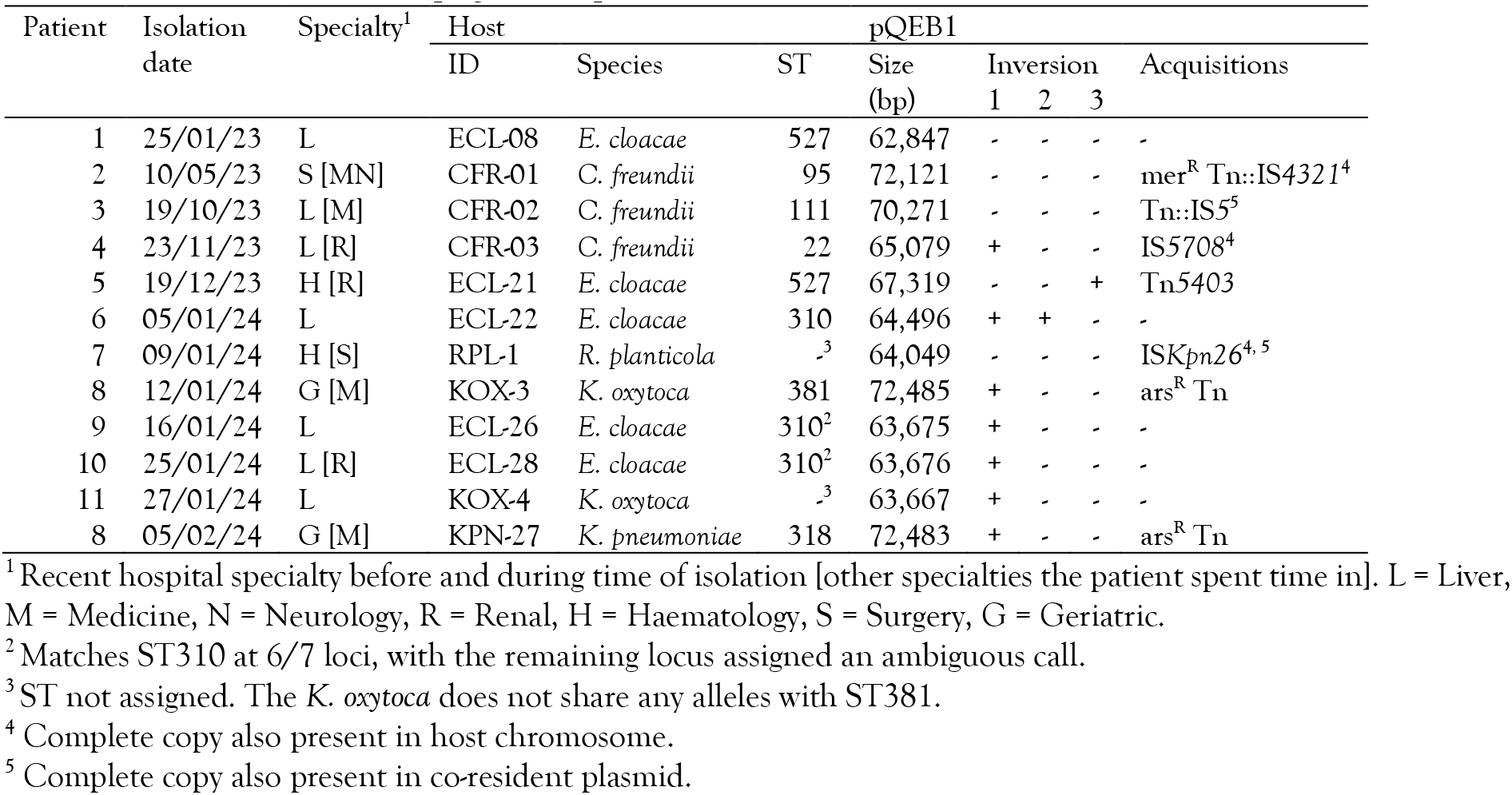
Characteristics of pQEB1 representatives and their bacterial hosts.

Most of the pQEB1 variants observed here were found in a single host each. However, variants derived from IS*26* inversion event 1 (pQEB_inv1) were found in five different bacterial hosts and six patients between November 2023 and February 2024. As expected for an IS*26*-mediated event ^11^, pQEB_inv1 inverted a 36.6 kb segment of the plasmid, including backbone and resistance region, generating a new copy of IS*26* and an 8 bp target site duplication (Figure 1A). Variation subsequent to pQEB1_inv1 indicates that this sub-lineage has continued to evolve as it has transferred between hosts, undergoing another IS*26*-mediated inversion, or acquiring IS*5708* or an 8,804 bp arsenic resistance transposon (Figure 1C). Plasmids with identical structures that include the arsenic resistance transposon were found in *K. oxytoca* and *K. pneumoniae* that were isolated from the same patient on the Geriatric ward, two weeks apart (Table 1).

## Discussion

Our observations of pQEB1 in both clinical and non-clinical CPEs at the QE over the past year are consistent with the characteristics of a conjugative plasmid outbreak. Representatives of pQEB1 have been found in multiple bacterial hosts that were isolated from 11 patients who stayed in multiple wards across two buildings on the QE site. The presence of pQEB1 at UHCW in May 2023 is concordant with the QE outbreak, and we believe explained by the fact that the UHCW patient had recently returned to UHCW after spending time at the QE for a procedure. The observed distribution of pQEB1 over this 13-month period suggests that it has significant transmission potential in nosocomial settings. Although lineages related to it have been detected internationally ^9,12^, pQEB1 exhibits a distinct structure that has so far only been seen in isolates from the English West Midlands. In future, it will be interesting to track any further spread of the pQEB1 lineage within and beyond this region of the UK.

While the data presented here strongly suggests that pQEB1 is disseminating in this nosocomial setting, it cannot be used to determine the precise locations in which horizontal transfer is occurring, whether that be within patients or the hospital environment. Focused surveillance might reveal environmental reservoirs of carbapenem resistance plasmids. If so, environmental studies should assist with the identification of targets for targeted infection prevention and control (IPC) interventions that aim to reduce the spread and persistence of CRGs in hospitals. Nonetheless, our observation here that the pQEB1_inv1 variant was strongly associated with the QE Liver speciality, where it was found in multiple patients and bacterial hosts, suggests that this plasmid successfully persisted there. The QE Liver speciality is one of the largest in the UK, and provides a comprehensive range of hepatology, liver surgery and liver transplantation services. The nature of this unit requires frequent, high-level use of broad-spectrum antibiotics including meropenem and piperacillin-tazobactam, as well as ciprofloxacin and co-amoxiclav, shaping conditions that seem likely to select for successful carbapenem resistance plasmid lineages.

In addition to the epidemiological insights, our approach has afforded us the opportunity to study the evolutionary biology of the pQEB1 lineage. We observed the emergence of several distinct plasmid sub-lineages through the actions of smaller mobile genetic elements. In most cases, copies of mobile elements that appeared to be newly-acquired by pQEB1 variants were found in the chromosome or co-resident plasmids of their bacterial hosts at time of isolation. This suggests that, given the extent observed in this relatively small sample set, there might be frequent exchange of smaller genetic elements between plasmids and chromosomes as plasmids disperse horizontally and pass through new hosts. Concerningly, the pQEB1 lineage already carries copies of IS*26*, and we have observed IS*26* activity *in situ* in the form of inversion events.

IS*26* is a major driver of resistance gene accumulation in Gram-negative bacteria, and carriage of it predisposes plasmids to acquiring further accessory gene-bearing IS*26* TUs ^13^. Though the acquisition of new TUs was not observed here, pQEB1 is primed for the accumulation of further antibiotic resistance genes, or for the formation of cointegrates with other plasmids, as has been observed for a different IS*26*-associated N-type plasmid lineage that was found in a *Proteus mirabilis* in China ^4^.

Using only sequence data generated with Oxford Nanopore R10.4.1 flow cells has proven to be a cost- and time-efficient method for conducting surveillance of CPEs at the QE. Generating complete plasmid sequences has facilitated an examination of structural variation, which has provided insights into the epidemiology and evolutionary biology of the pQEB1 lineage. We expect this approach will be a useful addition to hospital IPC strategies, and will continue to further our understanding of mobile genetic elements contributing to the emergence of extensive and pan-antibiotic resistance in hospitals globally.

## Methods

Bacterial strains were grown in LB broth (VWR Chemicals) at 37°C with 180 rpm shaking overnight. Genomic DNA was extracted from overnight cultures using the Monarch Genomic DNA Purification Kit (New England Biolabs) and quantified using the Qubit Broad Range dsDNA kit (ThermoFisher). Genomic DNA was prepared for sequencing with the SQK-NBD114 barcoding kit and sequenced on R10.4.1 flow cells using the GridION platform (Oxford Nanopore Technologies), with the addition of bovine serum albumin as recommended by the manufacturer. Reads were assembled using hybracter ^14^. Plasmid replicons, insertion sequences and antibiotic resistance genes were detected using PlasmidFinder, ISFinder and ResFinder, respectively ^15–17^. MLST was performed using mlst (https://github.com/tseemann/mlst). Complete plasmid sequences were visualised and annotated manually using Gene Construction Kit (Textco Biosoftware).

## Data availability

Sequencing reads are available from NCBI under BioProject Accession PRJNA1106791. The complete sequences of all pQEB1 structural variants described here are in Supplementary File 1.

